# A systemic role of macrophage-derived BMP2/4 homolog Dpp in regulating sterol hormone synthesis under dietary stress

**DOI:** 10.1101/2025.09.10.675304

**Authors:** Sergio Juarez-Carreño, Marco Milán

**Affiliations:** Institute for Research in Biomedicine (IRB Barcelona), The Barcelona Institute of Science and Technology, Baldiri Reixac, 10, 08028 Barcelona, Spain; Institució Catalana de Recerca i Estudis Avançats (ICREA), Pg. Lluís Companys 23, 08010 Barcelona, Spain

## Abstract

A high-sugar diet (HSD), prevalent in Western countries, compromises the growth and development of children and primes their bodies for chronic diseases. Macrophages, which are immune cells found throughout the body, can sense environmental changes and influence surrounding tissues. Using *Drosophila* as a model, here we show that macrophages respond to an HSD by increasing the production of Decapentaplegic (Dpp), a protein related to human BMP2/4. Dpp then signals to the endocrine system, the prothoracic gland, to inhibit the synthesis of ecdysone, a crucial steroid hormone that triggers the transition from larva to pupa. By reducing ecdysone synthesis, the larval stage is prolonged. This extended developmental period provides larvae exposed to an HSD more time to reach an optimal size. Our findings highlight how different organs communicate to counteract the negative impact of an HSD during development and expand our understanding of Dpp function, showing that it acts not only as a local developmental signal but also as a systemic regulator.

## Introduction

Western dietary patterns are commonly characterized by high intakes of refined sugars, processed foods, and saturated fats^1^. The low production costs of high-sugar foods contribute to their disproportionate prevalence among socioeconomically disadvantaged populations, thereby exacerbating existing health disparities^2^. This dietary paradigm is strongly correlated with an increased susceptibility to numerous chronic diseases, including Type 2 Diabetes Mellitus, insulin resistance, obesity, cardiovascular pathologies, neurological disorders, chronic systemic inflammation, and adverse developmental outcomes^3,4^. Of particular concern is the observed phenomenon of delayed pubertal onset (developmental timing) in obese children^5,6^, potentially attributable to the impaired synthesis of crucial puberty-driving steroid hormones^7^. Experimental animal models subjected to high-calorie dietary interventions consistently recapitulate the aforementioned human pathologies, demonstrating detrimental effects on development, such as pubertal delay, systemic inflammation, and insulin resistance^8–10^. The nutritional modulation of systemic physiology is facilitated by the secretion of signaling molecules from specialized sensing organs, including adipose tissue^11,12^, the gastrointestinal tract^13^, and macrophages^14^. These inter-organ communication pathways are critical for orchestrating physiological responses to external stimuli and maintaining organismal homeostasis^15,16^. Notably, *Drosophila* larvae subjected to a high-sugar diet (HSD) show a sugar-dose-dependent delay in the larval-to-pupal transition^17^, an analogous developmental process to mammalian puberty, alongside chronic inflammatory states^18^. Furthermore, recent research highlights the capacity of macrophages to perceive dietary alterations and modulate, as a response, the physiological state of the tissues in which they reside^14,19,20^. Specifically in *Drosophila*, macrophages are also involved in sensing periods of inanition, thereby influencing ecdysone synthesis—the sole steroid hormone in insects—during development, which in turn influences developmental timing^21^. In this study, we shed light on the mechanistic role of macrophages in regulating ecdysone synthesis under an HSD. The combined use of transcriptional profiling of macrophages subjected to an HSD and functional genetic analyses has allowed us to unravel the role of macrophage-derived Decapentaplegic (Dpp) in inhibiting ecdysone synthesis and delaying the larval-to-pupal transition. We present experimental evidence that the extended developmental time gives larvae exposed to an HSD more time to reach an optimal size. These results underscore a pivotal role of macrophages in the regulation of steroid hormone synthesis under high-calorie dietary regimens, thus positioning these cells as potential therapeutic targets for the management of metabolic disorders impacting on development.

## Results

### A high-sugar diet induces developmental delay and Dpp expression in macrophages

An HSD is defined as a nutritional condition characterized by a significantly elevated sugar concentration intake relative to a standard diet. Notably, HSDs have been used in *Drosophila* to mimic the physiological and developmental alterations observed in humans, including insulin resistance and chronic inflammation^8,18,22^. Larvae growing under a 10x HSD (4% sucrose in control diet vs. 40% sucrose in HSD, HSD larvae) showed a ∼24-hour (h) developmental delay compared to those growing in control or standard dietary conditions (control larvae) at the onset of metamorphosis, also known as the larval-to-pupal transition (Fig. 1A). Exposure to an HSD also led to a smaller pupal body size (Fig. 1B). The observed negative effect on systemic growth reflects signs of insulin resistance and resembles the phenotype of mutations in the insulin receptor or insulin-like peptides^23,24^. To explore whether the observed developmental delay is caused by the inhibited production of ecdysone—a hormone synthesized in the neuroendocrine prothoracic gland (PG) that drives the larval-to-pupal transition—we measured the levels of enzymes involved in its synthesis^25^, namely *neverland* (*nvl*), *phantom* (*phm*), and *shadow* (*sad*), at 104 h after egg laying (AEL), when the onset of pupariation takes place in control larvae. Notably, the mRNA levels of *nvl*, *phm*, and *sad* at 104 h AEL were significantly reduced in HSD larvae when compared to control conditions (Fig. 1C). Consistently, the expression of the ecdysone signaling target gene *eip75b* (*e75b*)^26^ was also reduced in HSD larvae (Fig. 1D). Macrophages are ancillary cells distributed throughout the body and they can sense environmental changes and impact on surrounding tissues by supplying growth factors and cytokines^16,27,28^. For example, macrophages in mammals are able to respond to high-sugar and fat diets and are involved in the pathophysiology of obesity and insulin resistance in adipose tissue^29^. Similarly, *Drosophila* macrophages can detect periods of inanition and modulate ecdysone synthesis and developmental timing^21^. Given these considerations, we addressed whether macrophages also play a role in sensing the dietary stress caused by an HSD and whether they contribute to the regulation of ecdysone synthesis and the resulting developmental delay. To this end, we used the pan-hemocyte driver *hemolectinΔ* (*hmlΔ*) to label differentiated macrophages with the fluorescence protein GFP (*hmlΔ>gfp*). Under an HSD, the number of macrophages was reduced compared to controls (Fig. 1E), suggesting that macrophages respond to dietary stress. To further characterize this response, we carried out bulk RNA-sequencing of FACS-sorted macrophages of larvae subjected to an HSD and compared their transcriptional profile to macrophages of control larvae. Broadly, Gene Ontology (GO) analysis indicated that HSD macrophages showed lower mRNA levels of genes related to carbohydrate catabolism and biosynthesis (Fig. S1A). GO analysis also pointed to an upregulation of lipid metabolism (Fig. S1A and S1B), suggesting that macrophage function relies on triglycerides as an energy source, as has been postulated in the context of HSDs^30^. Moreover, under an HSD, macrophages surprisingly reoriented towards homeostatic functions instead of classical immunological ones, a finding supported by the observed upregulation of growth factor-related GO terms (Fig. S1A, and S1B). Among the secreted proteins whose mRNA levels were significantly increased in HSD macrophages, we found imaginal disc growth factors (*idgf1*, *idgf2*, *idgf3*, *idgf4*), the TNFα homolog *egr*, PDGF/VEGF growth factor *pvf2*, the BMP homolog *dpp*, the FGF homolog *pyr*, and the growth-blocking factor *gbp3* (Fig. 1F). These observations point to a potential role of macrophages in sustaining homeostatic functions through inter-organ communication with distal organs. Of these secreted molecules, our attention was drawn to Dpp, which has been previously shown to modulate BMP signaling in the PG and to negatively regulate ecdysone production when overexpressed in distant tissues^31,32^. The expression levels of Brinker (Brk), a transcriptional repressor negatively regulated by Dpp, were reduced in the macrophages of HSD larvae (Fig. 1F). The increase in *dpp* levels was confirmed by RT-qPCR, where we used two housekeeping genes *Rp49* and *Act5C* to avoid the potential dysregulation of metabolic genes by the HSD (Fig. 1G). We next analyzed whether BMP signaling was activated in the PG of these larvae. The phosphorylation of the BMP transcription factor Mothers of Dpp (Mad)^33^ and the absence of Brk expression were used to monitor the activation of BMP signaling^34–36^. pMad levels were significantly increased (Fig. 1F) and Brk protein levels decreased (Fig. 1G) in the PG of HSD larvae (Fig. 1G). All these observations point to a potential role of macrophage-derived Dpp in driving the developmental delay caused by the HSD.

**Figure 1.**
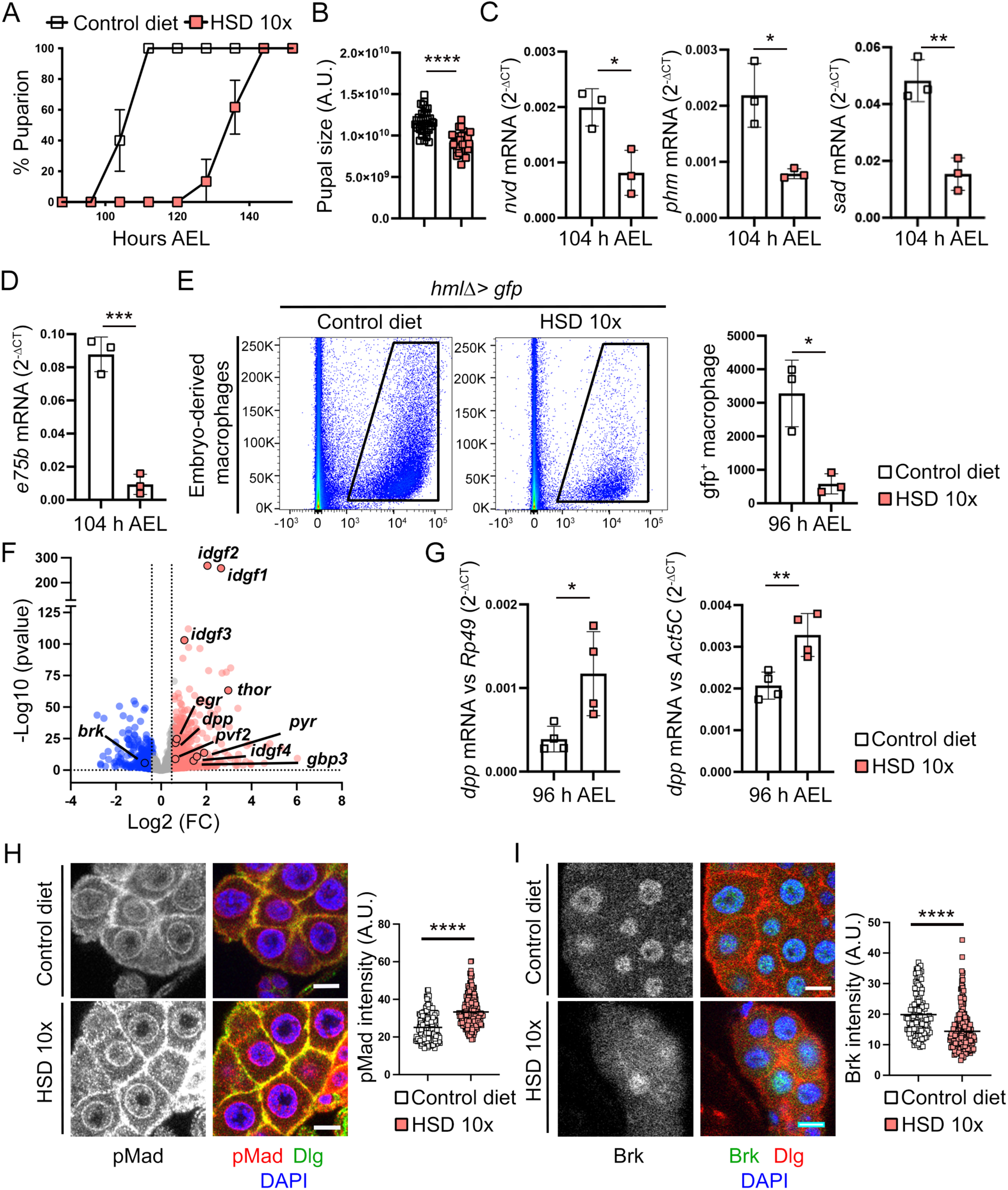
A high-sugar diet induces developmental delay and a transcriptional response in macrophages. (A, B) Developmental timing (A) and pupal size (B) of larvae growing in control diet or a 10x high-sugar diet (HSD) (A: n=3, 20 larvae per experiment in three independent crosses, 60 larvae in total per condition; B: n≥34). (C, D) Transcriptional levels of the ecdysone biosynthesis enzymes *nvl*, *phm* and *sad* (C) and the ecdysone signaling target gene *e75b* (D) in larvae growing in control diet or HSD. Samples were pooled by five larvae per condition. mRNA levels were normalized to *rp49*. n=4. (E) Flow cytometry and counts of total GFP-positive macrophages (*hmlΔ>gfp*) of larvae growing in control diet and HSD. n=3. (F) Volcano plot illustrating the differential gene expression in macrophages. These macrophages were isolated from larvae exposed to either a control diet or HSD (n=4). Red indicates upregulation and blue indicates downregulation of gene expression under the HSD condition (adjusted P<0.05; |fold change|≥1.5). Relevant growth factor genes are labeled with their corresponding gene symbols. Fold changes and p-values were calculated using the DESeq2 R package. (G) Transcriptional levels of *dpp* in larvae growing under control diet and HSD. mRNA levels were normalized to *rp49* and *Act5C*. Samples were pooled by five larvae per condition. n=4. (H, I) pMad (H) and Brk (I) protein expression levels in prothoracic glands of 96 h AEL-old larvae growing in control diet and HSD. Images show a Z-stack and scale bars of 10 μm. n≥6. n≥6. Values represent the means ± SD. Data were analyzed by two-tailed unpaired t test, and. *P < 0.05, **P < 0.01, ***P < 0.001, and ****P < 0.0001.

### Macrophage-derived Dpp regulates ecdysone synthesis and developmental timing under an HSD

To test whether macrophage-derived Dpp can regulate developmental timing and ecdysone synthesis under an HSD, we combined the use of *hmlΔ-Gal4* driver and a *dpp-RNAi* to deplete *dpp* expression specifically in macrophages (Fig. S2A). Pupariation in control larvae took place at ∼104 h AEL, whereas HSD larvae pupariated 24 h later (at ∼136h AEL). The developmental delay caused by the HSD was partially rescued by macrophage-specific depletion of *dpp* (Fig. 2A). Interestingly, the developmental delay caused by the diet was reproduced by overexpressing *dpp* in macrophages of control larvae (Fig. S2B). These data reveal a pivotal role of macrophage-derived Dpp in delaying developmental timing. To explore whether Dpp regulates developmental timing by inhibiting ecdysone synthesis in the PG, we next studied the mRNA levels of ecdysone synthesis enzymes *nvl*, *phm*, and *sad*, as well as the ecdysone signaling target gene *e75b* in HSD and control larvae. The increase in the mRNA levels of these three enzymes took place at ∼104 h AEL in control larvae but was delayed by 24 h in HSD larvae and was much milder (Fig. 2B). Interestingly, macrophage-specific depletion of *dpp* partially restored the peak mRNA levels of *nvl*, *phm*, and *sad* in HSD larvae, but this increase was still delayed by 24 h compared to control larvae (Fig. 2B). This observation is consistent with the partial impact of macrophage-specific depletion of *dpp* on developmental timing (Fig. 2A). Similar effects on the expression levels of the ecdysone target gene *e75b* were observed in HSD-larvae upon macrophage-specific depletion of *dpp* (Fig. 2C). To corroborate the inhibitory role of macrophage-derived Dpp in ecdysone synthesis, we next analyzed the total levels of ecdysteroid hormones in HSD and control larvae upon *dpp* depletion. Again, in HSD larvae, the increase in the ecdysone levels that precedes the onset of metamorphosis was delayed by 24 h and was much milder (Fig. 2D). Macrophage-specific depletion of *dpp* also restored the increase in the ecdysone levels, but this increase still occurred 24 h later than in controls (Fig. 2B). Taken together, these data support the notion that macrophage-derived Dpp inhibits ecdysone synthesis under HSD conditions, thereby modulating developmental timing. We did not observe macrophages near to or in close contact with the PG under either physiological or HSD conditions (Fig. S2D). To explore whether macrophage-derived Dpp can travel from distant macrophages to the PG, we used a transgenic fly expressing a functional Dpp::GFP fusion protein^37^. We searched for the presence of GFP fluorescence in the PG when Dpp::GFP was specifically expressed in macrophages. Interestingly, the GFP signal was observed in the PG of these larvae, indicating that macrophage-derived Dpp is secreted and interacts with PG cells (Fig. 2E). Taken together, these findings indicate that macrophage-derived Dpp contributes to the HSD-driven developmental delay through inhibition of ecdysone synthesis.

**Figure 2.**
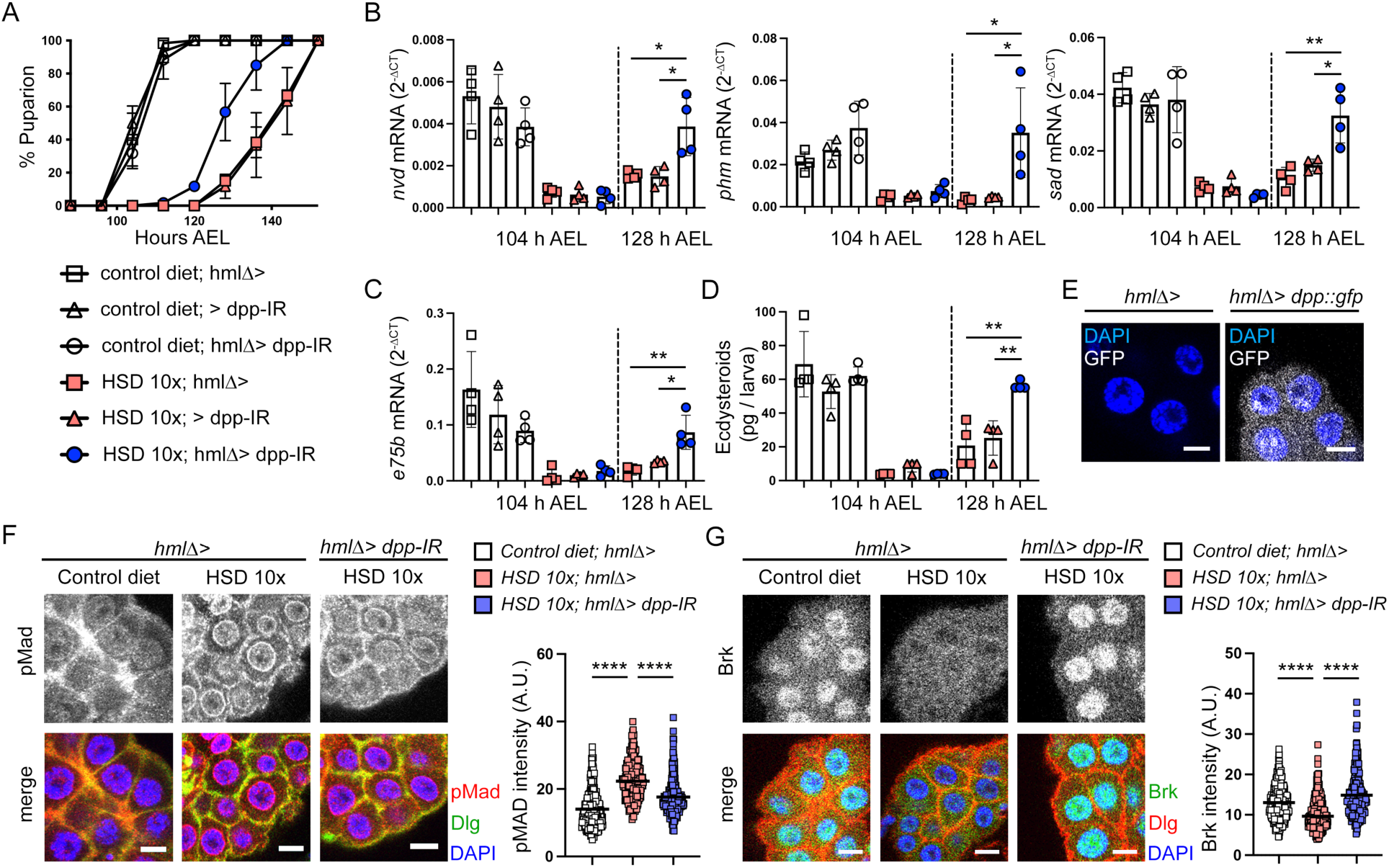
Macrophage-derived Dpp signals to the prothoracic gland to inhibit ecdysone synthesis and to cause a developmental delay under high-sugar diet. (A) Developmental timing in larvae of the indicated genotypes growing in control diet or HSD (n=3, 20 larvae per genotype in three independent crosses, 60 larvae in total per condition and genotype). (B, C) Transcriptional levels of the ecdysone biosynthesis enzymes *nvl*, *phm*, and *sad* (B) and the ecdysone signaling target gene, *e75b* (C) were quantified across the indicated conditions, genotypes and time points. mRNA levels were normalized to *rp49*. n=4. (D) Total ecdysteroid levels in the indicated genotypes and conditions and quantified by ELISA at the indicated time points. n = 4, with each sample pooled from 10 larvae. (E) Macrophages can secrete a Dpp tagged with GFP, which can be detected in the prothoracic gland. n≥5. (F, G) pMad (F) and Brk (G) protein expression levels in the prothoracic gland at 96 h AEL in the indicated genotypes and conditions. Images in E-G show a Z-stack and scale bars of 10 μm. F: n≥7. G: n≥6. Values represent the means ± SD. Data in B, C, and D were analyzed by two-tailed unpaired t test, and *P < 0.05, and **P < 0.01. Data in F and G were analyzed by non-parametric Kruskal–Wallis test and multiple comparison and ****P < 0.0001.

### Macrophage-derived Dpp signals to the PG to regulate ecdysone synthesis and developmental timing under HSD conditions

The increase in pMad levels observed in the PGs of HSD larvae was rescued upon macrophage-specific depletion of *dpp* (Fig. 2F), as was the reduction in the levels of Brk (Fig. 2G). These data support a role of macrophage-derived Dpp in activating BMP signaling in the PG. We next assessed the functional role of BMP signaling in the PG in regulating ecdysone production and the developmental delay in HSD larvae. To this end, we combined the use of a PG-specific Gal4-driver *(phm-GAL4)* and *RNAi* to knock down the Dpp receptor Thickveins (Tkv) and the BMP signaling transcription factor Mad and analyze the impact of the HSD on developmental timing. PG-specific depletion of *tkv* fully reverted the HSD-induced increase in pMad levels and the reduction in Brk levels in the PG (Figure 3A, B). Interestingly, PG depletion of *tkv* and *mad* partially rescued the developmental delay caused by the HSD (Fig. 3C and S3A). Similar results on developmental timing were obtained upon expression of a dominant negative isoform of Tkv (Tkv^DN^) in the PG (Fig. S3B)^38^. These data support the hypothesis that macrophage-derived Dpp delays developmental timing through Tkv and Mad in the PG under HSD conditions. To explore the role of Tkv and Mad in regulating ecdysone production, we measured the mRNA levels of ecdysone synthesis enzymes *nvl*, *phm*, and *sad*, as well as the ecdysone signaling target gene *e75b* in HSD larvae upon PG-specific depletion of *tkv* and *mad*. The increase in the *nvl*, *phm*, and *sad* mRNA levels that occurred in control larvae at ∼104 h AEL was delayed by 24 h in HSD larvae, and this increase was much milder (Fig. 3D and S3C). Interestingly, PG-specific depletion of *tkv* or *mad* partially restored the mRNA levels of *nvl*, *phm*, and *sad* in HSD larvae, but this increase was still delayed by 24 h (Fig. 3D and S3C). This observation is consistent with the partial impact of the PG-specific depletion of *tkv* or *mad* on developmental timing (Fig. 3C and S3A). Similar effects on the expression levels of the ecdysone target gene *e75b* were observed in HSD-larvae upon PG-specific depletion of *tkv* or *mad* (Fig. 3E and S3D). To further confirm the role of BMP signaling in regulating ecdysone synthesis in the PG, we compared the total levels of ecdysteroid hormones in HSD and control larvae upon PG-specific depletion of *tkv*. The increase in the ecdysone levels that precedes the onset of metamorphosis in control larvae was delayed by 24 h and was much milder in HSD larvae (Fig. 3F). In this regard, PG-specific depletion of *tkv* restored the increase in ecdysone levels, but this increase still occurred 24 h later than in controls (Fig. 3F). All these results indicate that macrophage-derived Dpp acts on the PG to inhibit, through Tkv (Fig. 3G) and Mad, ecdysone synthesis and developmental timing in larvae subjected to HSD conditions.

**Figure 3.**
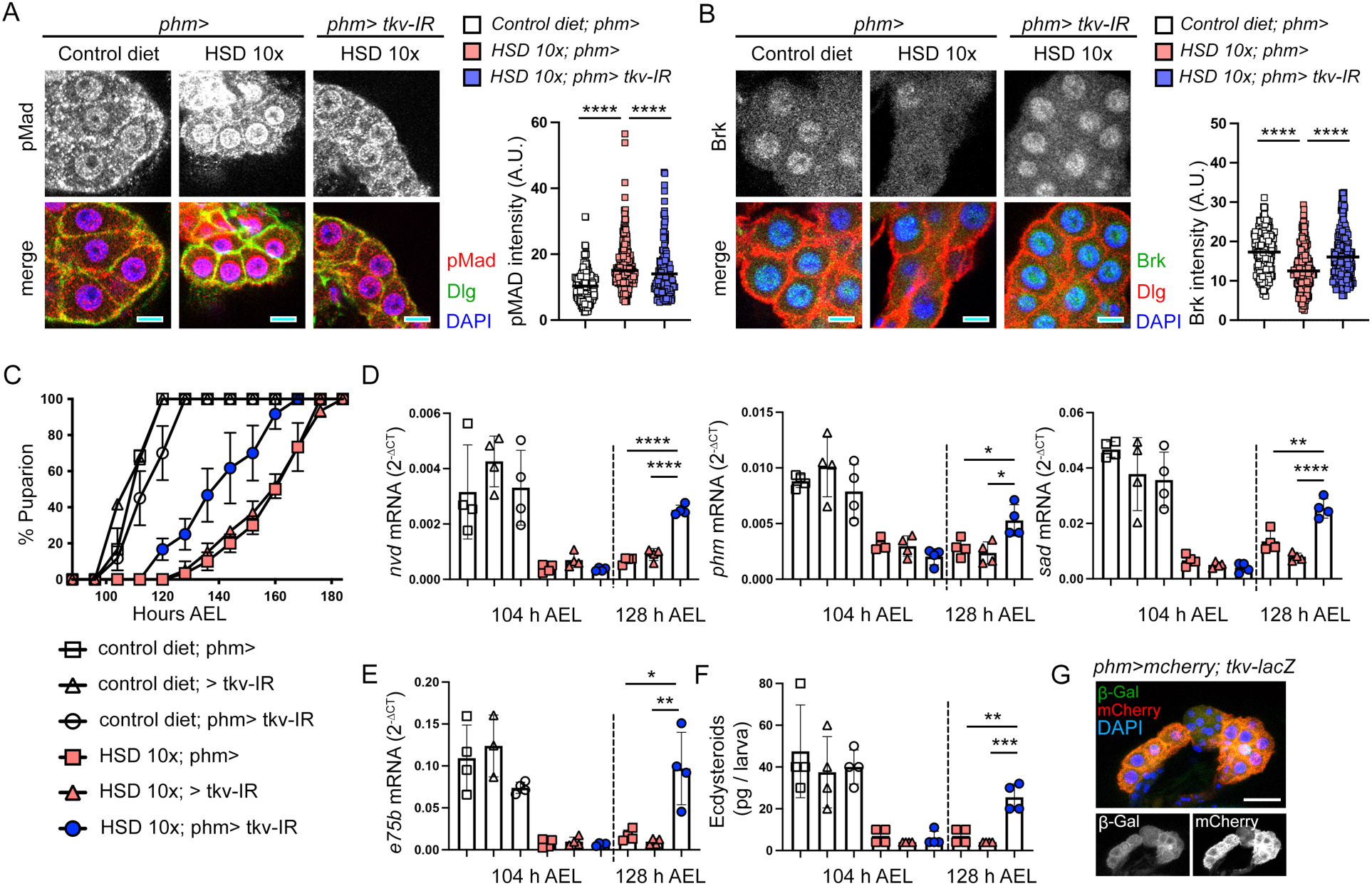
Dpp signaling in the prothoracic gland inhibits ecdysone synthesis and induces a developmental delay under a high-sugar diet. (A, B) pMad (A) and Brk (B) protein expression levels in the prothoracic gland at 96 h AEL in the indicated genotypes and conditions. n≥5. (C) Developmental timing in larvae of the indicated genotypes growing in control diet or HSD (n=3, 20 larvae per genotype in three independent crosses, 60 larvae in total per condition). (D, E) Transcriptional levels of the ecdysone biosynthesis enzymes *nvl*, *phm*, and *sad* (D) and the ecdysone signaling target gene, *e75b* (E) measured across the indicated conditions, genotypes and time points. mRNA levels normalized to *rp49*. n=4. (F) Total ecdysteroid levels in the indicated genotypes and conditions, quantified by ELISA at the indicated time points. n = 4, with each sample pooled from 10 larvae. (G) Tkv expression in the prothoracic gland. Images in A, B, G show a Z-projection and scale bars of 10 μm (A, B) or 30 μm (G). Values represent the means ± SD. Data were analyzed in D, E, and F by two-tailed unpaired t test, and *P < 0.05, **P < 0.01, ***P < 0.001, and ****P < 0.0001. Data were analyzed in A and B by non-parametric Kruskal– Wallis test and multiple comparison and ****P < 0.0001.

### Dpp-driven developmental delay dampens the negative effects of an HSD on systemic growth

It is well-established that ecdysone can regulate growth in two different ways. First, during early stages of larval development, ecdysone is present at low levels and interacts with its receptor, EcR, in somatic tissues to promote proliferative growth^39^. Second, the ecdysone peak that occurs at the larval-pupal transition modulates the overall growth period of larvae^40^. Thus, when this peak is delayed, larvae have more time to grow and display a larger pupal size^41^. Conversely, when the peak is induced prematurely, larvae have less time to grow and show a smaller pupal size^42^. In the context of an HSD, the observed reduction in body size is generally thought to be a consequence of the detrimental effects of the diet on insulin resistance^8^. We thus questioned whether the effects of an HSD on extending developmental timing via the production of Dpp by circulating macrophages serve to minimize the impact of insulin resistance on body growth, allowing the animal to reach a minimal size necessary for adult biological functions. To test this hypothesis, we analyzed whether partial restoration of developmental timing was able to enhance the negative impact of the HSD on pupal size. Interestingly, macrophage-specific depletion of *dpp*, which partially rescued the developmental delay caused by the HSD (Fig. 2A), further increased the reduction in pupal size caused by this dietary regimen (Fig. 4A). Similar effects on developmental timing and pupal size were obtained upon PG-specific depletion of *tkv* or *mad* (Fig. 4B, C) or expression of *tkv^DN^* (Fig. S3E). We observed that macrophage-specific depletion of *dpp* or PG-specific depletion of *tkv* or *mad* in control larvae did not have an impact on pupal size (Fig. 4A). Interestingly, *dpp* overexpression in the macrophages of control larvae reproduced the effects of ecdysone inhibition in delaying developmental timing (Fig. S2B) and increasing pupal size (Fig. S2C). Overall, these data support the notion that the modulation of developmental timing by macrophage-derived Dpp under HSD conditions serves as a buffering mechanism to allow the organism to extend its developmental period, thereby enabling it to reach a minimum final size sufficient for adult functions.

**Figure 4.**
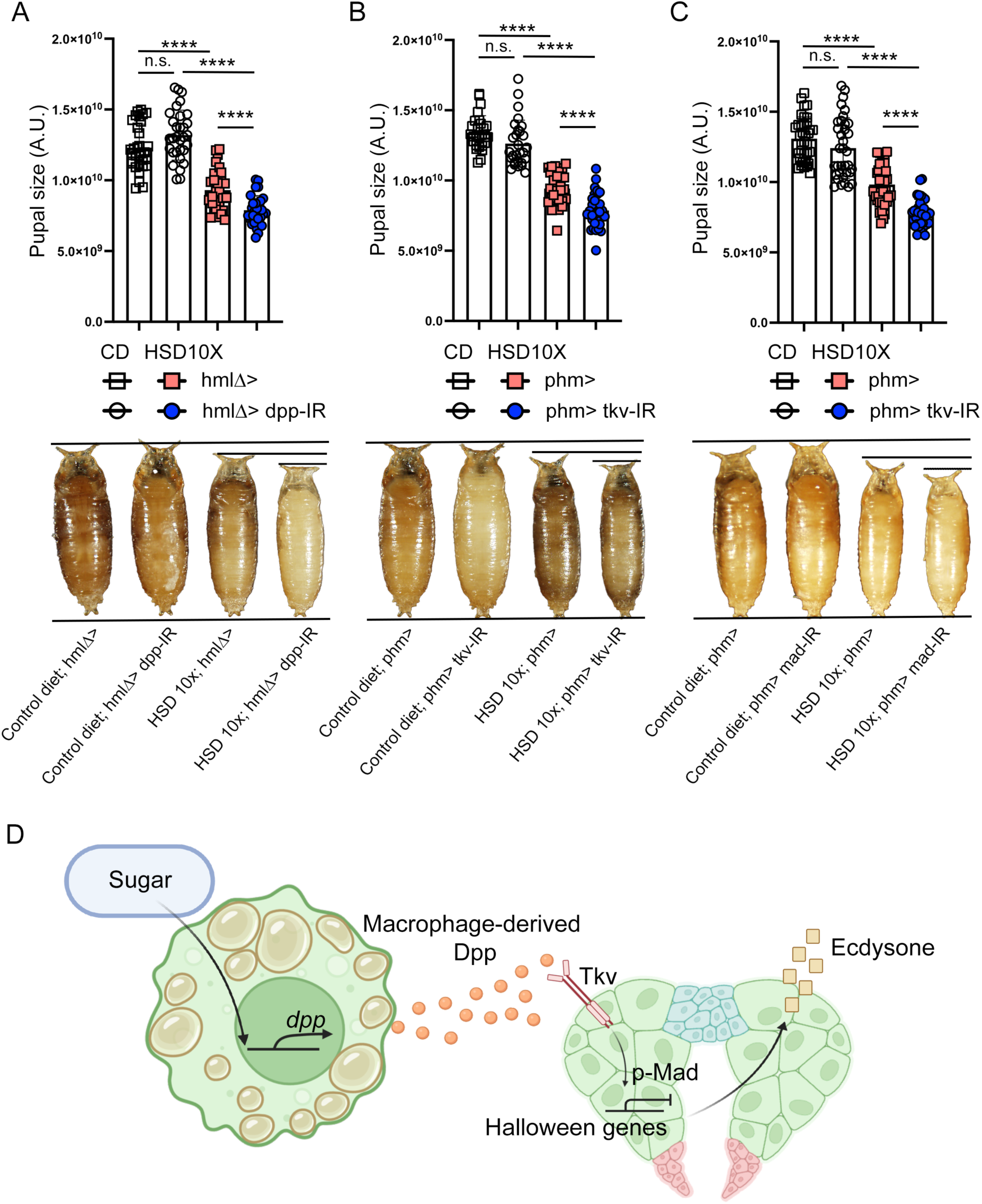
Macrophage-derived Dpp signals to the prothoracic gland to regulate systemic growth under dietary stress. (A-C) Final pupal size of larvae subjected to macrophage-specific depletion of *dpp* (A), *tkv* (B) and *mad* (C), and growing in control diet (CD) or HSD. A: n≥30, B: n≥28. C: n≥35. A.U., arbitrary units. Values represent the means ± SD. Data were analyzed by two-tailed unpaired t test and. ****P < 0.0001. (D) Macrophage-secreted Dpp regulates ecdysone biosynthesis and developmental progression under conditions of diet-induced stress, primarily through its receptor Thickveins (Tkv) expressed in the prothoracic gland. Scheme created with BioRender.com.

## Discussion

Developmental timing, including the onset of puberty, is genetically determined and intricately mediated by the biosynthesis of sterol hormones^43–47^. However, this finely tuned developmental program can be significantly modulated by external factors such as tumors and nutritional status^48–52^. Recent research has underscored the crucial role of macrophages as pivotal sensor cells capable of detecting environmental challenges and modulating the physiology of distal organs to maintain systemic homeostasis^14,16,53–56^. Specifically in mammals, macrophages have been postulated to stimulate the production of sexual sterol hormones within the gonads^57,58^. In *Drosophila*, the macrophage genetic cassette *inr/dtor/pvf2* modulates sterol hormone biosynthesis under periods of inanition during development^21^. Here, we demonstrate that macrophages also regulate ecdysone synthesis in *Drosophila* under HSD conditions. Specifically, macrophages secrete the BMP 2/4 homolog factor Dpp to inhibit ecdysone biosynthesis under these dietary conditions, thereby extending developmental timing and allowing the organism to reach an optimal body size. Our findings indicate that Dpp interacts with the Tkv receptor in the PG, thereby activating BMP signaling and consequently inhibiting the expression of ecdysone biosynthesis enzymes (Fig. 4D). Here we elucidate the specific physiological conditions and anatomical source from which Dpp is secreted and acts as an inter-organ signal to mediate ecdysone synthesis inhibition in the neuroendocrine organ, beyond its established roles as a morphogen in the imaginal disc^59,60^. Furthermore, this study highlights the critical role of macrophages, particularly macrophage-secreted BMPs, in mediating the progression of pathological conditions such as diet-induced insulin resistance. Notably, BMP factors and signaling have previously been implicated in promoting insulin resistance and obesity in mice, and their dysregulation improves insulin sensitivity^61–63^. Additionally, these findings have potential implications for understanding the regulation of sterol hormones across the spectrum of diabetes and insulin resistance in mammals^64–66^, offering mechanistic insights into phenomena observed in obese children, where puberty is often delayed in those with diabetes^67,68^.

## Materials and methods

### Drosophila husbandry

The plasmatocyte-specific reporter line and driver *hmlΔ-gal4;UAS-2xeGFP* stock was a gift from S. Sinenko^69^. Prothoracic gland-specific driver *phm-gal4* was a gift from Michael O’Connor^42^, *UAS-tkv^DN^* was a gift from Ernesto Sánchez-Herrero, *UAS-dpp::gfp* was a gift from Aurelio Telemann. *dpp-TRiP.JF02455-RNAi* (36779), *tkv-TRiP.HMS02185-RNAi* (40937), *mad-TRiP.JF01264-RNAi* (31316), *tkv-lacZ* (11191), *20xUAS-6xmCherry-HA* (52268), and *hmlΔ-gal4* (30139) lines were from the Bloomington Stock Center at Indiana University (Bloomington, IN). Flies were reared in standard medium (4% glucose, 55 g/L yeast, 0.65% agar, 28 g/L wheat flour, 4 ml/L propionic acid and 1.1 g/L nipagin) at 25°C on a 12-h:12-h light:dark cycle. For all experiments, larvae of mixed sexes were used.

### Standard diet and 10x high-sugar diet

Strains of *Drosophila melanogaster* and control experiments (control diet) were fed standard medium (4% glucose, 55 g/L yeast, 0.65% agar, 28 g/L wheat flour, 4 ml/L propionic acid and 1.1 g/L nipagin) at 25°C in 12:12h light-dark cycles. For the high-sugar diet, we added 10x sucrose to the standard food, increasing the total sugar content to 40%.

### Egg-laying and pupariation formation measurements

We crossed 20 males with 20 females. After 48 h, the adult flies were transferred to grape agar plates (Nutri-fly grape agar premix, Genesee Scientific) supplemented with yeast paste. They were allowed to lay eggs for 3 to 4 h at 25°C. Following egg deposition, the adults were removed, and the eggs were incubated at 25°C for an additional 48 h. At this point, second-instar larvae were collected and transferred to either standard *Drosophila* food (20 larvae per tube) to maintain physiological conditions or to a 10x high-sugar diet to induce nutritional stress. All larvae were reared at 25°C. We monitored the larval-to-pupal transition every 8 h, with “time 0” defined as 4 h after the onset of egg laying, referred to as “AEL” (after egg laying).

### Pupae volume measurements

Twenty males and twenty females were crossed and allowed to lay eggs over a 24-h period. The adult flies were transferred to fresh tubes every 24 h, and the deposited eggs were incubated at 25°C. Resulting pupae were collected and photographed with their dorsal side facing upward. Measurements of length and width were taken using ImageJ, and pupal volume was calculated using the following formula: v = 4/3π(L/2)(l/2)2 (L, length; and l, width)^70^. A Leica MZ216F microscope was used to measure pupal volume, and an Olympus MVX10 macroscope for representative pupae imaging.

### Immunohistochemistry

The brain–ring gland complex was dissected out in cold phosphate-buffered saline (PBS) buffer. It was then fixed in 4% formaldehyde for 20 min^71,72^ and stained with antibodies in PBS with 0.3% BSA and 0.2% Triton X-100. Next, the brain–ring gland complex was stained overnight at 4°C with goat anti-GFP (ab6673) (1/500; abcam), mouse anti-Dlg (4F3) (1/50; Hybridoma bank), rabbit anti-Smad3 (phospho S423 + S425) (C25A9) or pMAD (1/100; Cell Signaling Technology), chicken β-Gal (ab9361) (1/500; abcam), and guinea pig anti-Brinker (1/100; a gift from Gines Morata). Secondary antibodies, Fluorescein (FITC) AffiniPure Donkey Anti-Goat IgG (H+L) (1/250) (RRID: AB_2340400), Alexa fluor 488-conjugated AffiniPure Donkey Anti-Mouse IgG (H+L) (1/250) (RRID: AB_2341099), Alexa Fluor 488 AffiniPure Donkey Anti-Chicken IgY (IgG) (H+L) (1/250) (RRID: AB_2340375), Cy3-AffiniPure Donkey Anti-Mouse IgG (H+L) (1/250) (RRID: AB_2315777), Cy5 AffiniPure Donkey Anti-Guinea Pig IgG (H+L) (1/250) (RRID: AB_2340462), and Cy3 AffiniPure Donkey Anti-Rabbit IgG (H+L) (1/250) (AB_2307443) were purchased from Jackson ImmunoResearch. For DNA and nuclei staining, we used DAPI (1/1000). We set up the tile scan with the micrometer-thick Z stack on a Zeiss LSM880 microscope with a 40× oil objective. Images were acquired with a resolution of 1024 × 1024.

### Confocal microscopy

Samples were analyzed using a Zeiss LSM 780 confocal microscope. Image acquisition was done with a 40× oil objective. The most representative image is shown for each experiment.

### Flow cytometry analysis of macrophage and macrophage counts

Hemocytes from five larvae at 96 h AEL were bled into 120 μl of FACS buffer (PBS, 0.5% bovine serum albumin, and 2 mM EDTA) containing 1 nM phenylthiourea (Sigma-Aldrich, St. Louis, MO, USA) to prevent melanization. We used two forceps to make an incision from the posterior and pull to the anterior ends of the larvae. We allowed larvae to bleed for a few seconds, and then carefully scraped the lateral cuticle to avoid the lymph gland disruption^73^. Macrophages (GFP-positive hemocytes and 40,6-diamidino-2-phenylindole (DAPI) negative] were extracted from *hmlΔ>2xeGFP* larvae and quantified on a BD FACSARIA III (BD Biosciences). Cells were selected based on their size with an FSC-A from 2.5K to 250K and their granularity with an SSC-A from 2.5K to 150K. Later, we gated for singlets (FSC-H vs. FSC-A). For fluorescent hemocytes, we used a BlueC (530_30) RB-A laser (GFP). Flowcytometry plots were obtained with FlowJo 9.9.

### qRT-PCR on sorted macrophages and whole larvae

For qRT-PCR, 15,000 hemocytes were sorted into 600 μl of TRIzol LS (Thermo Fisher Scientific: 10296010). RNA extraction was performed following the manufacturer’s instructions (RNeasy Mini Kit, Qiagen). RNA concentration was measured with a SpectraMax QuickDrop UV-Vis Spectrophotometer from Molecular Devices. cDNA preparation was performed with the Invitrogen SuperScript IV First-Strand Synthesis System (Thermo Fisher Scientific) as per the manufacturer’s instructions. qRT-PCR was done with 5 to 10 ng of cDNA.

For whole larvae, qRT-PCR tissue homogenization was performed using a pestle on four to five larvae per sample, and RNA extraction was performed following the manufacturer’s instructions (RNeasy Mini Kit, Qiagen). For whole larvae, the same protocol as for the sorted cells was used, and the qRT-PCR was performed on 500 ng of cDNA. qRT-PCR was performed using QuantStudio6 Flex Real-Time PCR from Life Technologies and using TaqMan Fast Advance Mas-termix and TaqMan probes for *rp49* (Dm02151827_g1), *phm* (Dm01844265_g1), *nvd* (Dm03419116_m1), *sad* (Dm02139323_g1), *eip75b* (Dm01793666_m1), and *dpp* (Dm01842959_m1).

### Ecdysteroid measurement

For ecdysteroid extraction, 10 whole larvae from each condition (from 104 h AEL and 128 h AEL, as indicated in the figure) were collected in 2 ml of tissue-homogenizing mixed beads and then frozen in liquid nitrogen and transferred to −80°C. Samples were homogenized using a tissue lyser in 0.3 ml of methanol. They were then centrifuged at maximum speed for 5 min at room temperature, and the supernatant was transferred to another Eppendorf tube. We repeated this procedure by adding 0.3 ml of methanol to the pellet and mixing using vortex, and a third round using 0.3 ml of ethanol. The pooled sample after homogenization was 0.9 ml per sample. For quantification, the extracted samples were centrifuged at maximum speed for 5 min (room temperature) to remove any remaining debris and then divided into two tubes to generate technical replicates. The cleared samples were evaporated overnight^74^. The following steps were performed using the 20-HE enzyme-linked immunosorbent assay kit instructions (Bertin Pharma, no. A05120; 96 wells). Absorbance was measured at 410 nm with a microplate reader (BioTek Synergy H1 Plate Reader).

### RNA purification, library preparation, and bulk RNA-sequencing

Total RNA from a total of 4 replicates of 20,000 GFP^+^ macrophages was then purified at the IRB Barcelona Functional Genomics Core Facility, as described in González et al., 2010. mRNA was isolated from total RNA using the kit NEBNext Poly(A) mRNA Magnetic Isolation Module (New England Biolabs). NGS libraries were prepared from the purified mRNA using the NEBNext Ultra II RNA Library Prep Kit for Illumina (New England Biolabs). Eighteen cycles of PCR amplification were applied to all libraries. The final libraries were quantified using the Qubit dsDNA HS assay (Invitrogen) and quality-controlled with the Bioanalyzer 2100 DNA HS assay (Agilent). An equimolar pool was prepared with the 20 libraries and submitted for sequencing at the Centro Nacional de Análisis Genómico (CNAG). A final quality control by qPCR was performed by the sequencing provider before paired-end 150 nt sequencing on a NovaSeq6000 S4 (Illumina). More than 275 Gbp of reads were produced, with a minimum of 39 million paired-end reads per sample.

### -Data processing

Adapters from reads were removed using trim galore (v.0.6.7) (https://github.com/FelixKrueger/TrimGalore) with default parameters. Resulting reads were aligned to the dm6 *D. melanogaster* genome version using STAR (v.2.7.10a) (Dobin et al. 2013). The count matrix was generated in R (https://www.R-project.org/) through the function “featureCounts” of the Rsubread package (Liao, Smyth, and Shi 2019).

### ***-***Differential expression

Differential expression was measured using the DESeq2 package (Love, Huber, and Anders 2014). For plotting purposes and generating principal components, the count matrix was normalized using the “vst” function. The replicate batch was included in the model as a covariate.

### ***-***Gene set enrichment analysis

Gene set enrichment analysis was performed using the roastgsa package (Caballé-Mestres, Berenguer-Llergo, and Stephan-Otto Attolini 2023). GO and KEGG pathways were downloaded using the org.Dm.eg.db package.

### -Data availability

The raw and processed RNAseq data were deposited in the GEO database under accession number GSE303750. Reviewers can access the data at https://www.ncbi.nlm.nih.gov/geo/query/acc.cgi?acc=GSE303750 by entering the token mzgncqwalrkrlyf in the access box.

## Statistical analyses

All statistical analyses were performed using GraphPad Prism Software 9.0 with a 95% confidence limit (P < 0.05).

### Animal experiments

All animal experiments were done using *Drosophila melanogaster*, which does not require approval from the Ethics Committee.

## Acknowledgments

We thank the Bloomington Stock Center at Indiana University (Bloomington, IN) and the Developmental Studies Hybridoma Bank (USA) for providing the flies and antibodies. We are also indebted to the members of the Milán lab for helpful discussion on the manuscript and IRB Barcelona’s Advanced Digital Microscopy, Functional Genomics and Biostastics/Bioinformatics Facilities for their assistance. We gratefully acknowledge institutional funding from the Spanish Ministry of Science, Innovation and Universities through the Centres of Excellence Severo Ochoa Award, and from the CERCA Programme of the Catalan Government. M.M.’s lab is funded by a Spanish Ministry of Science, Innovation and Universities grant. S.J-C was awarded a Beatriu de Pinós Postdoctoral Fellowship, co-funded by the Marie Skłodowska-Curie program from Horizon 2020 and the Agency for Management of University and Research Grants of the Catalan Government.

## Author contributions

S.J-C. conceived the original idea, designed, supervised and performed all the experiments, contributed to funding acquisition and wrote the original manuscript. M.M. supervised the project, contributed to funding acquisition, and wrote the original manuscript.

## Data and materials availability

All data needed to evaluate the conclusions in the paper are present in the paper and/or the Supplementary Materials. Further information and requests for resources and reagents should be directed to and will be fulfilled by the Corresponding authors.

---

**Figure S1.**
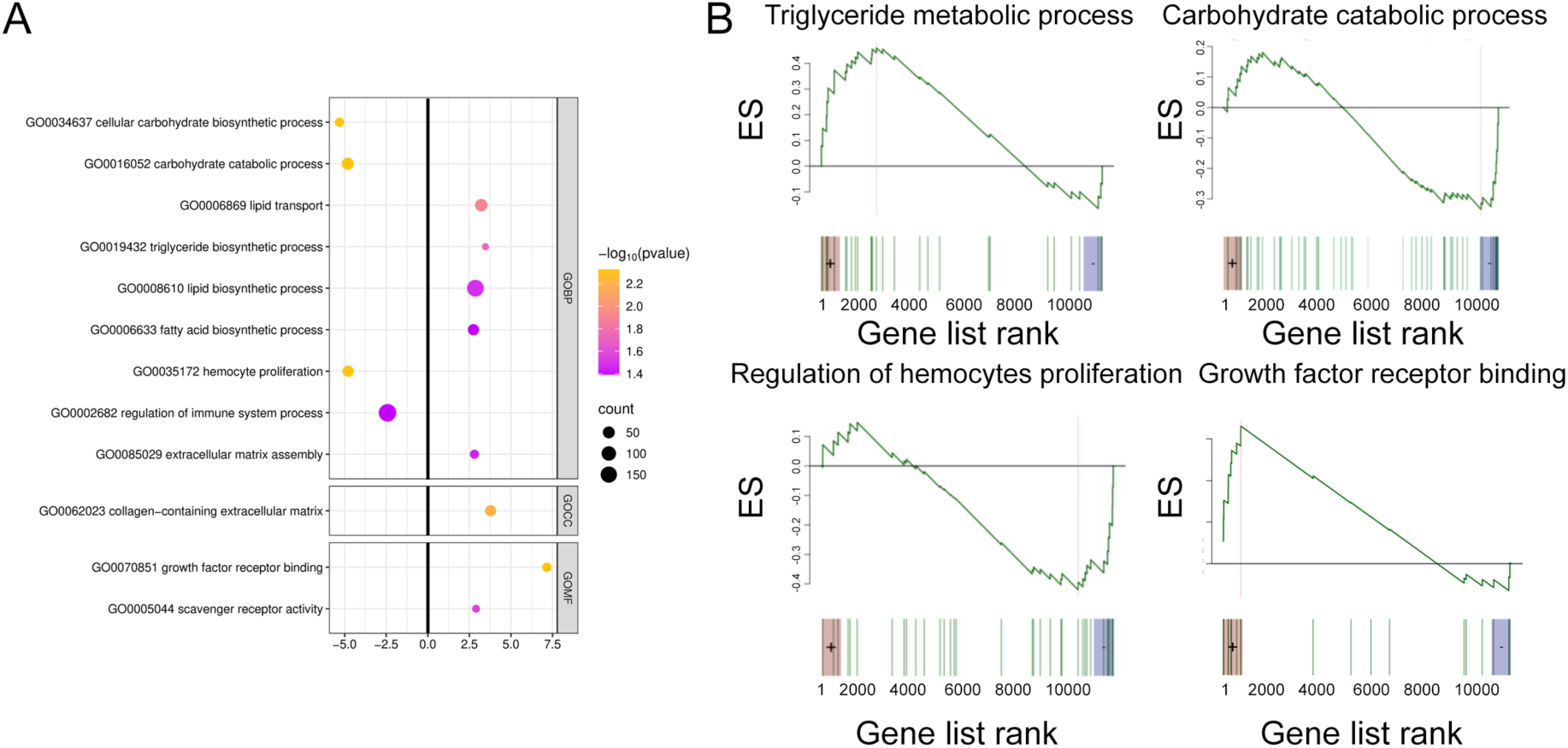
related to Fig. 1. Bulk RNA sequencing of macrophages shows metabolic switch and enhanced trophic role under nutrient-induced stress during development. (A) Gene Ontology (GO) enrichment results visualized using dot plots. Each dot represents an enriched GO term. The color scale indicates the direction and magnitude of gene expression changes, while dot size corresponds to the measured number of genes associated with each term. GO terms were considered significantly enriched at p value ≤ 0.05. Significance is visualized using −log₁₀ (p value), where higher values indicate stronger enrichment. (B) Gene set enrichment analysis of the GO pathways indicated.

**Figure S2.**
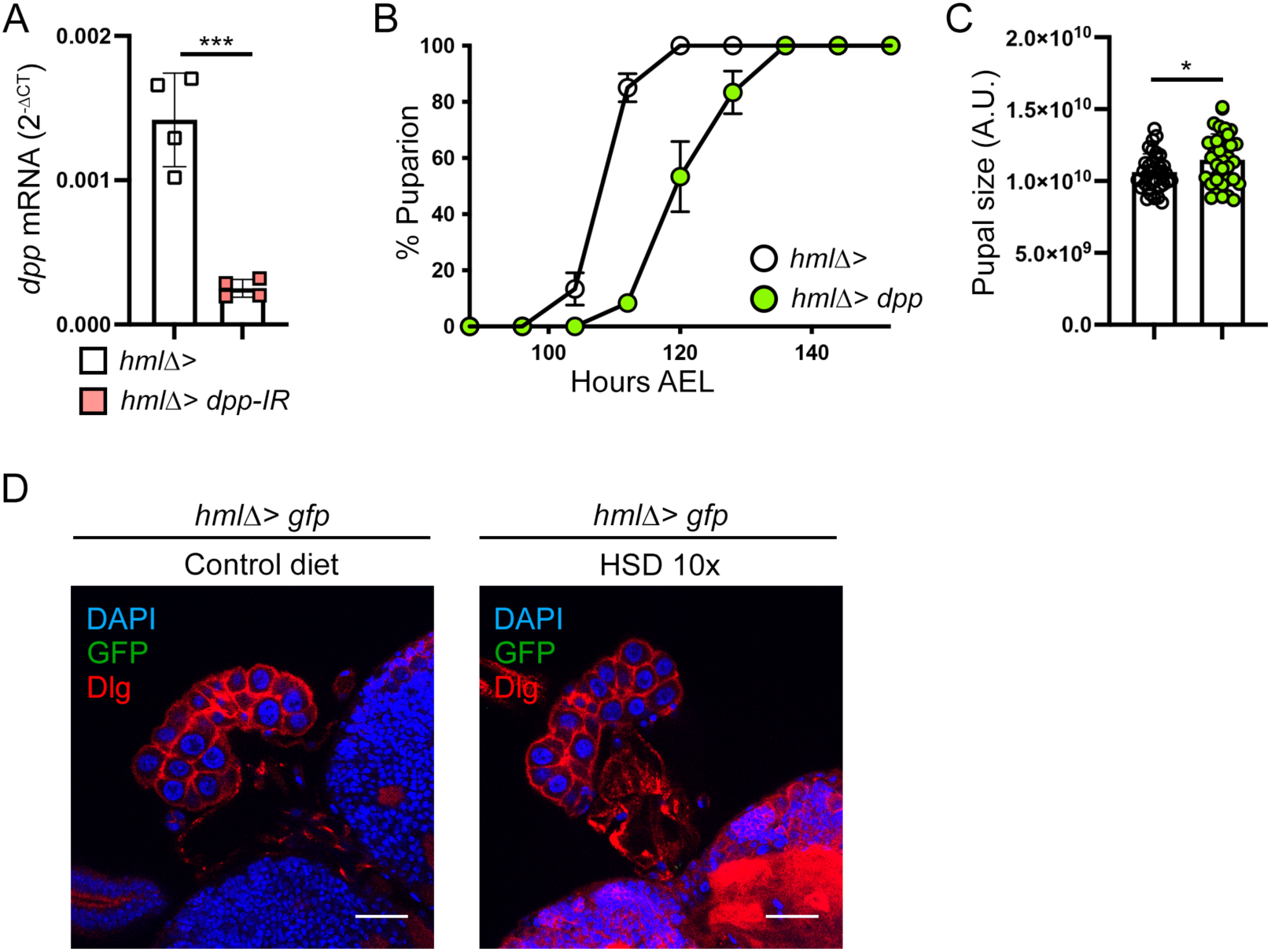
related to Fig. 2. Macrophages overexpressing Dpp induce a developmental delay and drive systemic growth. (A) RNAi efficiency of *dpp* RNAi line. mRNA levels were normalized to *rp49* at 96 h AEL of sorted macrophages in the indicated genotypes. n=4, with each sample pooled from 15,000 macrophages. (B) Developmental timing in larvae of the indicated genotypes and growing in control diet (n=3, 20 larvae per genotype in three independent crosses, 60 larvae in total per condition). (C) Final pupal size of larvae overexpressing *dpp* specifically in macrophages and growing in control diet. n=35. (D) Representative images of the prothoracic gland from larvae subjected to a physiological diet or 10x high-sugar diet. Macrophages are labeled with GFP fluorescent protein (*hmlτ1>gfp)*. Images show a Z-stack and scale bars of 30 μm. n=6. Data were analyzed by two-tailed unpaired t test, and values represent the means ± SD. *P < 0.05, and ***P < 0.001.

**Fig. S3.**
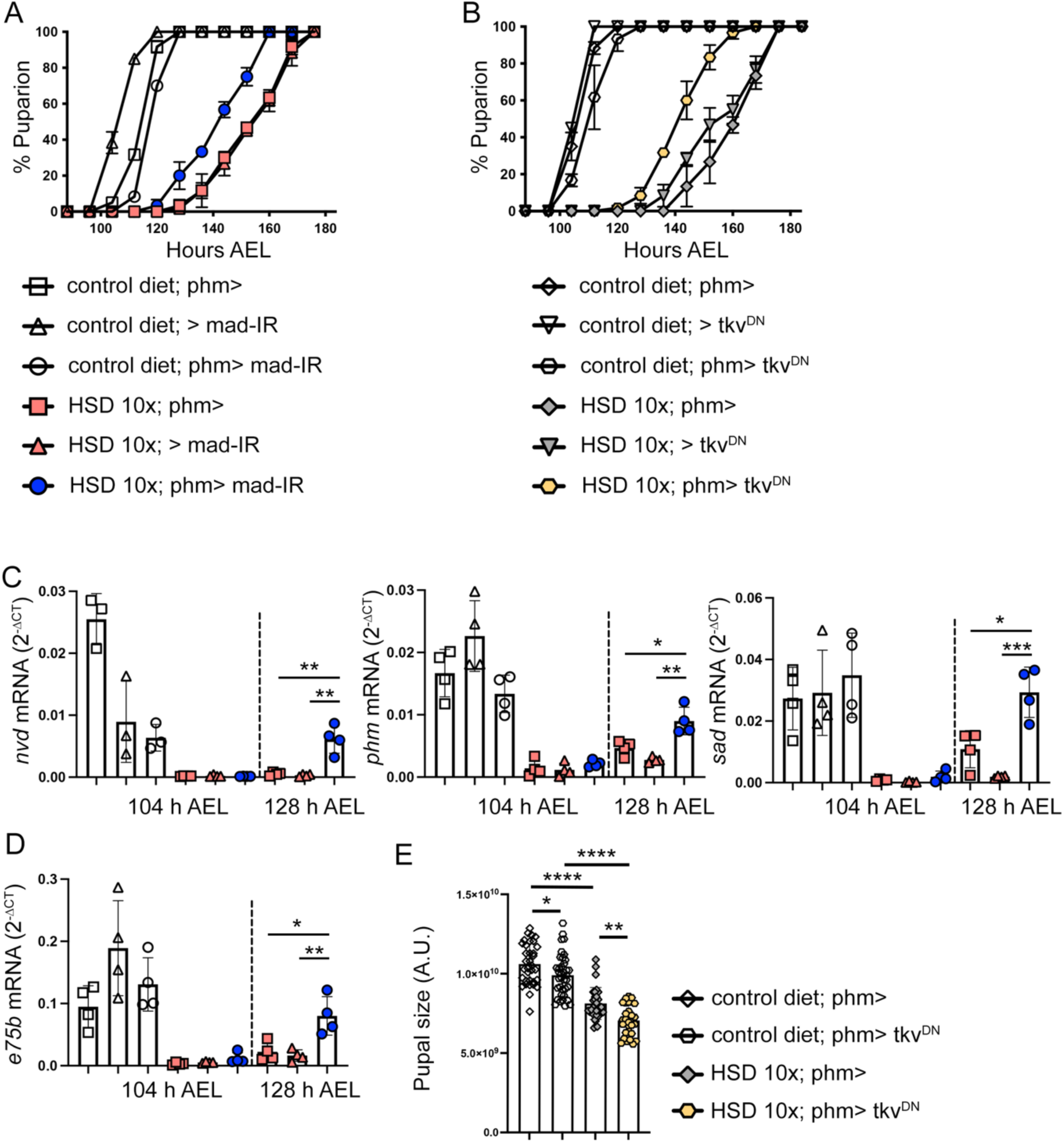
related to Fig.3 and Fig.4. A role of Dpp signaling in inhibiting ecdysone synthesis in the prothoracic gland and inducing a developmental delay under a high-sugar diet. (A, B) Developmental timing in larvae of the indicated genotypes and growing in control diet or HSD (n=3, 20 larvae per genotype in three independent crosses, 60 larvae in total per condition). (C, D) Transcriptional levels of the ecdysone biosynthesis enzymes, *nvl*, *phm*, and *sad* (C) and the ecdysone signaling target gene, *e75b* (D) were quantified across the indicated genotypes, conditions and time points. mRNA levels normalized to *rp49*. n=4. (E) Final pupal size of larvae growing in control diet or HSD and subjected to macrophage-specific expression of a dominant negative version of the Tkv receptor (Tkv-DN). n≥30. Data were analyzed by two-tailed unpaired t test, and values represent the means ± SD. *P < 0.05, **P < 0.01, ***P < 0.001, and ****P < 0.0001.

